# Stem nodulation: diversity and occurrence in *Aeschynomene* and *Sesbania* legumes from wetlands of Madagascar

**DOI:** 10.1101/2023.09.12.557367

**Authors:** Faustin F. Manantsoa, Marrino F. Rakotoarisoa, Clémence Chaintreuil, Adamson T.E. Razakatiana, Frédéric Gressent, Marjorie Pervent, Mickaël Bourge, Martial D. Andrianandrasana, Nico Nouwen, Herizo Randriambanona, Heriniaina Ramanankierana, Jean-François Arrighi

## Abstract

As an adaptation to flooding, few legume species have the original ability to develop nitrogen-fixing nodules on the stem. By surveying wetlands of Madagascar, we found a large occurrence and diversity of stem nodulation in Aeschynomene and Sesbania legumes. They represent opportunities to investigate different modalities of the nitrogen-fixing symbiosis in legumes.

## Introduction

The symbiosis between legume plants and soil rhizobia results in the formation of nitrogen-fixing nodules, generally exclusively appearing on the roots. However, in a handful of tropical legumes growing in wetlands, nodulation with rhizobia can also occur at stem-located dormant root primordia, a process that is referred as stem nodulation. Seen the semi-aquatic lifestyle of these legumes, it has been hypothesized that the stem nodulation trait is an evolutionary adaptation to flooding (Ladha *et al.*, 1992). Stem nodulation was first described for the African *Aeschynomene afraspera* (not to be confounded with *Aeschynomene aspera* found in Asia) (Hagerup, 1928). Stem nodulation gained agricultural interest after the discovery that the profuse stem nodulation as found in A. afraspera and Sesbania rostrata results in a high nitrogen fixation activity (Dreyfus and Dommergues, 1981; Alazard and Becker, 1987). To date, stem nodulation has been reported for species belonging to four legume genera: *Aeschynomene, Discolobium, Neptunia and Sesbania* (Boivin *et al.*, 1997). While, in the three latter genera, stem-nodulation has been described for one or very few species, more than 20 *Aeschynomene* species have been shown to form stem nodules (Chaintreuil *et al.*, 2013; 2016). These legume species can differ in their stem nodulation ability and intensity, and actually *S. rostrata* and *A. afraspera* are the ones for which stem-nodulation is the most profuse (Boivin *et al.* 1997a).

Strikingly, *S. rostrata* is stem-nodulated only by *Azorhizobium caulinodans* while other *Sesbania* species root-nodulate with rhizobia of the *Ensifer* genus (Boivin *et al.*, 1997a,b). Similarly, different types of *Bradyrhizobium* strains have been identified as nodulating *Aeschynomene species*, (Boivin *et al.*, 1997a; Alazard, 1985). The most important difference between the strains were the presence or absence of photosynthetic activity and nod genes to produce Nod factors (Molouba *et al.*, 1999; Giraud *et al.*, 2007; Miché *et al.*, 2011). So far, photosynthetic *Bradyrhizobium* strains have been exclusively found in nodules of stem-nodulating *Aeschynomene* species (Boivin *et al.*, 1997a) and strains lacking nod genes have been isolated from nodules of *Aeschynomene* species that cluster in a single clade (Chaintreuil *et al.*, 2013, 2016). The *Bradyhrizobium* ORS278-A. evenia interaction serves as model for the deciphering of the latter very specific interaction called Nod-independent symbiosis (Quilbé *et al.*, 2021). In contrast, strains having nod genes, such as *Bradyrhizobium* ORS285, have been isolated from nodules of *A. afraspera* that is one of the *Aeschynomene* species using a Nod-dependent interaction (Arrighi *et al.*, 2014; Brottier *et al.*, 2018). S. rostrata stem-nodulating *A. caulinodans* is not photosynthetic and has nod genes, but in both cases, the symbiotic interaction is very specific.

Although research has shed some light on the genetics of both partners in the *Bradyrhizobium-Aeschynomene* and *A. caulinodans-S. rostrata* symbiotic systems, our understanding of the molecular mechanism that causes stem nodulation is still in its infancy. Furthermore, our knowledge of the diversity and occurrence of stem nodulation *in natura* is relatively limited as it has been investigated in only a few geographical regions (e.g., James *et al.*, 2001; Molouba *et al.*, 1999; Miché *et al.*, 2011). To fill this gap, we conducted a field study in Madagascar that contains a variety of wetlands with an important plant biodiversity. A series of expeditions were organized to explore wetland-rich regions in the Central (RN1-Itasy Lake), Northern (Nosy Be), Western (RN4-Majunga) and Eastern (RN2-Alaotra Lake) parts of Madagascar (Fig. 1a). In these regions, we found stem-nodulated *Aeschynomene, Sesbania* and *Neptunia* spp. Ommiting these latters from this analysis, as nodules were only found on floating stems forming adventitious roots, a total of 69 Aeschynomene and *Sesbania* samples was collected (Table S1). Field observations were completed with molecular and flow cytometry analyses for accurate specimen identification (Tables S2 and S3). Here, we report on the stem-nodulated *Aeschynomene* and *Sesbania* species from Malagasy wetlands and discuss the opportunity of these resources to fuel research on the nitrogen-fixing symbiosis in legumes.

**Fig. 1.**
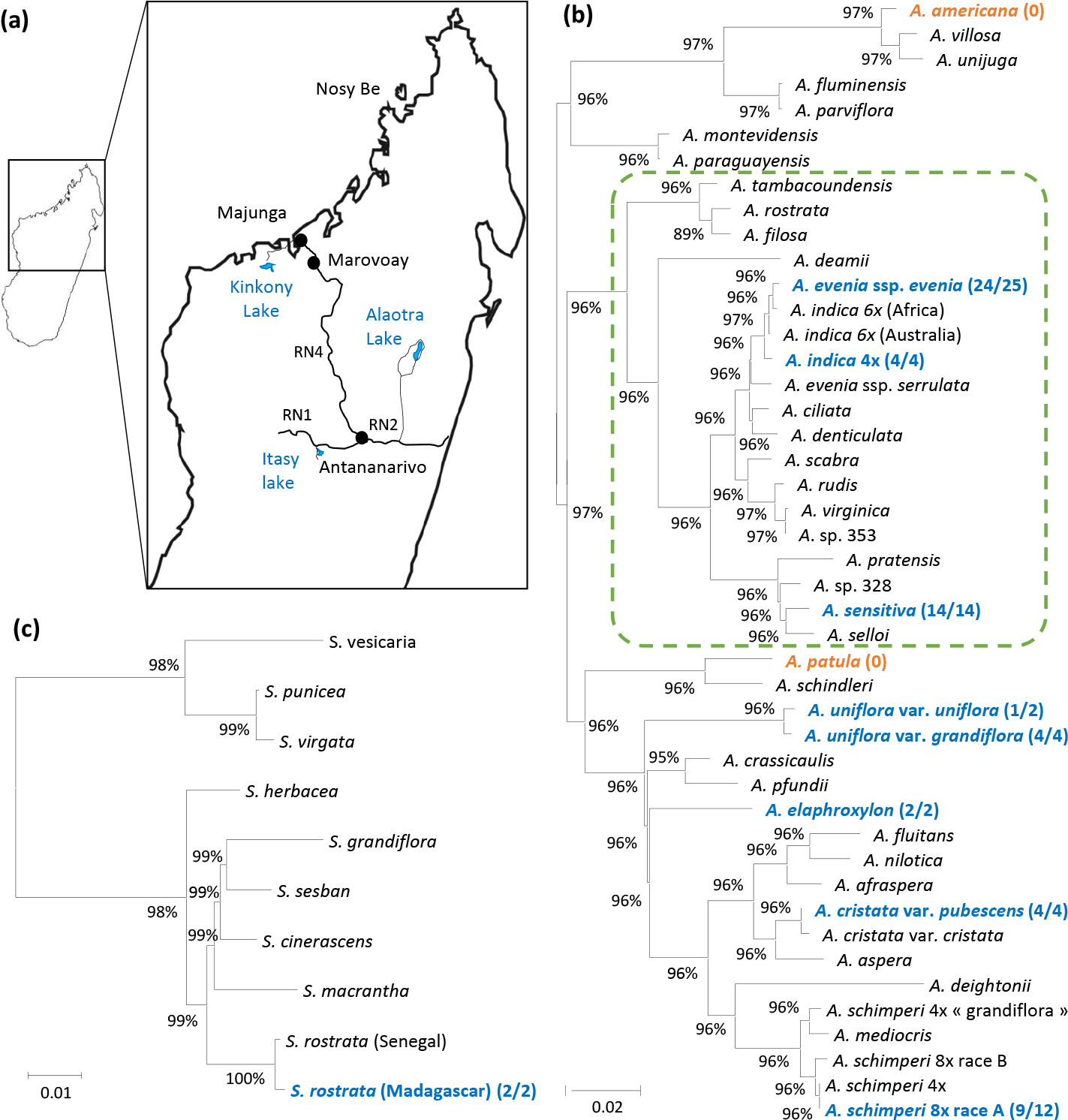
*Aeschynomene* and *Sesbania* species sampled in wetlands of Madagascar. (a) Map of Madagascar with a zoom on the four collecting sites, including RN1-Itsay Lake, RN2-Alaotra Lake, RN4-Majunga and Nosy Be. (b) Phylogeny of *Aeschynomene*. (c) Phylogeny of *Sesbania*. In (b) and (c), Maximum likelihood phylogenetic reconstructions were obtained using the concatenated the ITS + *matK* sequences. Numbers at nodes represent bootstrap values (% of 1000 replicates). Dashed box delineates the Nod-independent clade. Taxa collected in Madagascar are in bold and numbers on their right correspond to the occurrence of stem nodulation in the different sampling sites. In orange: no stem nodulation observed, in blue: stem nodulation observed in the present study.

## Stem nodulation in Nod-independent *Aeschynomene* species

Three stem-nodulated *Aeschynomene* species belonging to the Nod-independent clade were found in Malagasy wetlands (Fig. 1b). *A. evenia* was by far the most widespread one, being seen in all parts of Madagascar and in all wetland types: river banks, marshes and ricefields. *A. evenia* has a transatlantic distribution and a well-defined geographically-structured genetic diversity (Chaintreuil *et al.*, 2018). The form present in Madagascar was previously classified as the Eastern African genotype. As a result, it is closely related to the reference line from Malawi that was selected for the genetic dissection of the Nod-independent symbiosis (Quilbé *et al.*, 2021). Stem-nodulated *A. evenia* plants were observed in 24 out of the 25 sampling sites. Nodules were green, indicative of the presence of chloroplasts, and often located in the lower part of the stem, but their distribution could extend to the upper branch-containing part and they were usually present in a profuse fashion. These stem nodules were hemispherical and with a broad attachment to the stem (Fig. 2a).

**Fig. 2.**
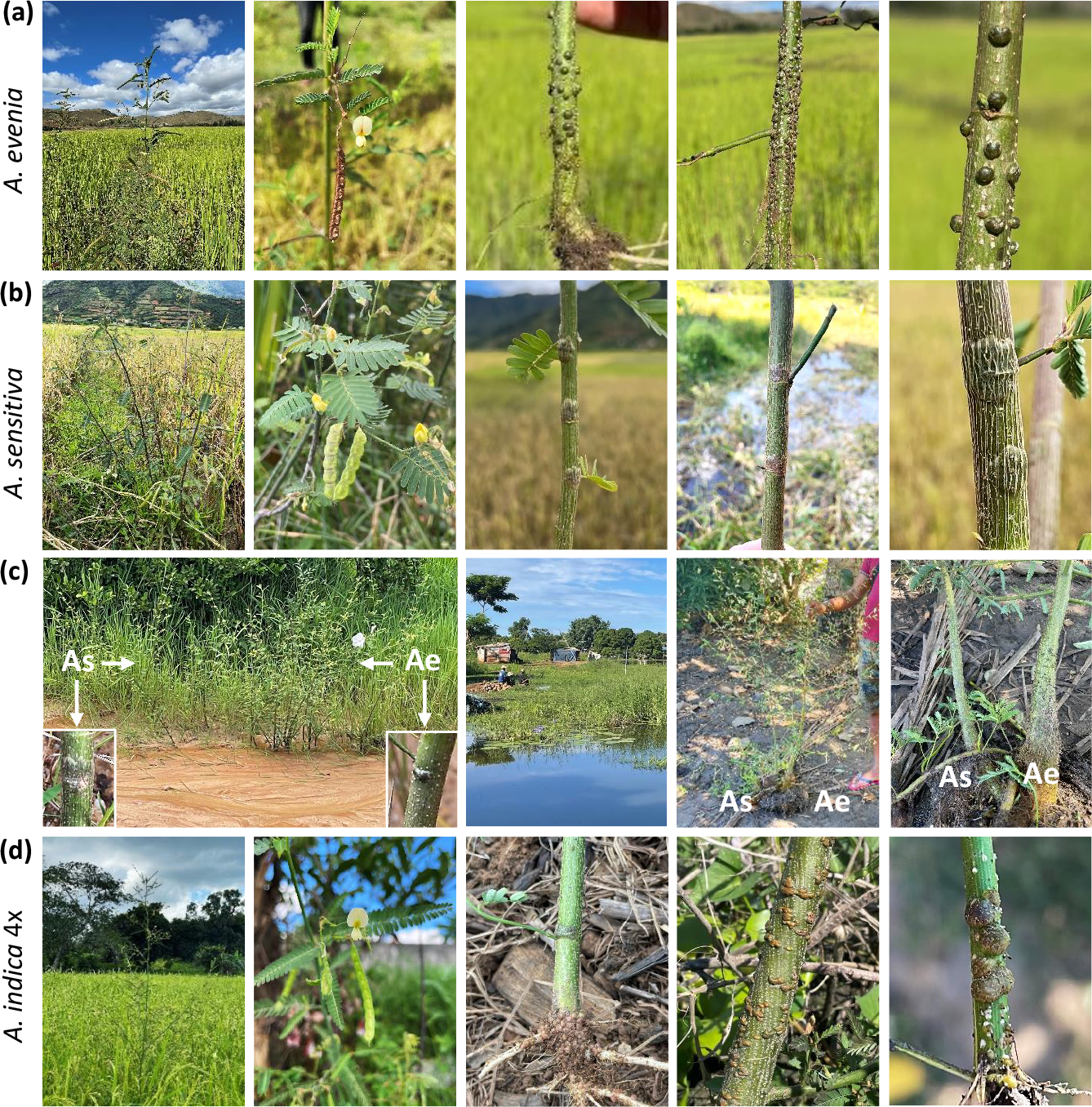
Stem nodulation in *Aeschynomene* species of the Nod-independent clade. (a) *A. evenia*. From left to right: 1-plant growing in a ricefield near Camp Bandro (Alaotra lake), 2-axillary axis bearing a yellow flower and a mature pod, 3-4-stem nodules located at the base or middle portion of the stem, 5-morphology of stem nodules. (b) *A. sensitiva*. From left to right: 1-plant growing in a ricefield at Manakambahiny, in the direction of Alaotra lake, 2-axillary axis bearing yellow flowers and developping pods, 3-4-more or less flattened stem nodules, 5-morphology of collar stem nodules. (c) Co-occurrence of *A. evenia* and *A. sensitiva*. From left to right: 1-Stand of mixed population in a shallow sandy river at Marofotroboka (on RN4), insets show stem nodules of *A. sensitiva* and *A. evenia*, respectively, 2-4-permanent marsh at Belobaka (Majunga) where *A. sensitiva* and *A. evenia* where found to grow side-by-side with entangled roots, both plants displaying stem nodules. Ae: *A. evenia*, As: *A. sensitiva*. (d) *A. indica* 4x. From left to right: 1-plant growing in a ricefield in Nosy Be, 2-axillary axis bearing a yellow flower and a developping pods, 3-4-stem nodules located near the base or at the middle portion of the stem, 5-morphology of stem nodules. Note the presence of numerous pink root nodules in 3.

The second typical species found in different wetlands is *A. sensitiva*. Similarly to *A. evenia*, this species has also a transatlantic distribution and the genotype occurring in Madagascar is concommitantly present in Africa and Eastern Brazil (Chaintreuil *et al.*, 2018). *A. sensitiva* is well-known because the model strain ORS278, a photosynthetic *nod* gene-lacking *Bradyrhizobium*, was isolated from stem nodules in Senegal (Giraud *et al.*, 2000, 2007). The photosynthetic activity of this strain was demonstrated to be important for the efficiency of stem nodulation. In addition, *A. sensitiva* has the particularity to develop unique ‘collar’ nodules on the stem (Fig. 2b). These were green and readily observed in all 14 sampling sites. Although *A. sensitiva* was globally less frequently found as compared to *A.evenia*, at the collection sites the two species were frequently growing adjacent to each other (Fig. 2c). The two species are known to have overlapping distributions in Madagascar but *A. sensitiva* is absent in the drier regions whereas *A. evenia* does (Dupuy *et al.*, 2002).

Unexpected was the discovery of *A. indica* in marshes from Majunga and in ricefields in Nosy Be because this species is not native of Madagascar. *A. evenia* and *A. indica* are morphologically similar but they can be distinguished from each other via the flowers that on *A. indica* plants are in general larger than those on *A. evenia* (Fig. 2d). This high resemblance is due to their belonging to a same polyploid species complex where A. evenia 2x, *A. indica* 4x and *A. indica* 6x forms are present (Arrighi *et al.*, 2014; Chaintreuil *et al.*, 2018). To confirm the visual species identification, we sequenced the nuclear ITS and chloroplastic *matK* gene and determined the genome size of the specimens JFA29 and JFA109. The obtained data were typical for A. indica 4x plants, confirming the visual identification (Fig. S1, Table S2) (Arrighi *et al.*, 2014). In the four collection sites, A. indica plants were stem-nodulated, the nodules varying in shape and size but always with an enlarged base and with a green color (Fig. 2d). Similar to othern *Aeschynomene* species, pink root nodules could be observed in unflooded conditions, (Fig. 2d).

## Stem nodulation in Nod-dependent *Aeschynomene* species

Among *Aeschynomene* species falling outside of the Nod-independent clade, the pantropical *A. americana* and the endemic *A. patula* were frequently present in explored sites. Both species are of interest because they have been proposed as model plants complementary to *A. evenia* to develop a comparative genetic system to study the Nod-independent and Nod-dependent symbioses in *Aeschynomene* (Brottier et al., 2018). However, for both species no stem-nodulated plants were found at the collection sites (Fig. 1b).

Two *Aeschynomene* species, *A. elaphroxylon* and *A. schimperi*, which are believed to be introduced in Madagascar, were found in the Central Plateaux (Fig. 1b) (Du Puy *et al.*, 2002). *A. elaphroxylon* was specially present around the Aloatra lake. It is a distinctive *Aeschynomene species* as it can form large shrubs to small trees with very showy yellow flowers and spiny stems (Fig. 3a). In the two sampled sites, plants were scarely nodulated and, if so, only on the submerged parts of the stems. In that case, nodules were green and had a flattened hemispherical shape with a broad attachment to the stem (Fig. 3a). In contrast, *A. schimperi* was frequently found in ricefields. For this species, previous genetic analysis uncovered the presence of 4x and 8x cytotypes, and Malagasy specimens correspond to the 8x cytotype (Chaintreuil *et al.*, 2016). In non-waterlogged conditions, numerous pink nodules could be observed on the main root, while under flooded conditions (9 of the 12 sampled sites) green nodules were present in the lower part of the stem (Fig. 3b). These nodules were spherical with a narrow attachment to the stem.

**Fig. 3.**
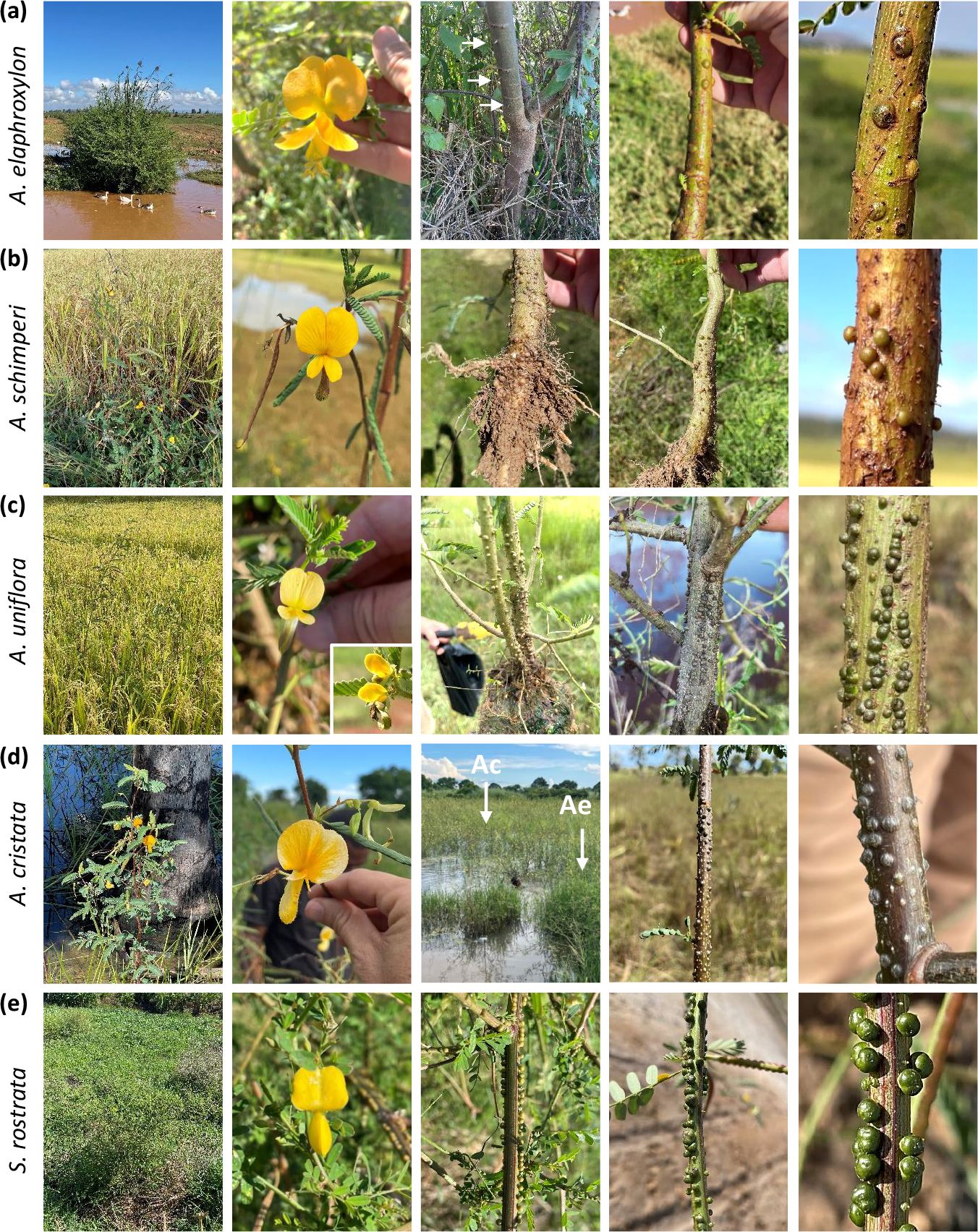
Stem nodulation in Nod-dependent *Aeschynomene* and *Sesbania* species. (a) *A. elaphroxylon*. From left to right: 1-stand of shrubby plants growing at the edge of a marsh near the Alaotra lake, 2-showy yellow flower with visible anthers, 3-woody stem with spines, 4-nodules developping at the base of the stem, 5-morphology of stem nodules. (b) *A. schimperi*. From left to right: 1-young plants in a ricefield at Maharefo, South of the Alaotra lake, 2-yellow flower, 3-pink root nodules, 4-stem nodules, 5-morphology of stem nodules. *A. uniflora*. From left to right: 1-plants growing in a ricefield at Bongomena on the RN4, 2-yellow flowers with the smaller variant form shown in the inset, 3 and 4-stem nodules of the of the normal and morphological variant, respectively, 5-morphology of stem nodules. (d) *A. cristata*. From left to right: 1-Plant growing in a swampy palm grove, 2-showy yellow flower, 3-Stand of *A. cristata* (Ac) in the center of a marsh lined with *A. evenia* (Ae), 4-nodules present all along the stem, 5-morphology of stem nodules. (e) *S. rostrata*. From left to right: 1-plant growing on the edge of a marsh at Amboromalandy on RN4, 2-Yellow flower, 3 and 4-arrows of nodules running all along on the stem, 5-morphology of stem nodules.

In regions at lower altitude, in the North and West of Madgascar, two other Nod-dependent *Aeschynomene* species were found. *A. uniflora* was present in the region of Majunga and in Nosy Be. Here it grew either at the edge of marshes or in ricefields where often also *A. evenia* or *A. sensitiva* species were present. Strikingly, two plant morphotypes were observed: one corresponding to erected plants with well-visible flowers (3 sampled sites), and a second represented by shrubby plants bearing small flowers (two sampled sites) (Fig. 3c). It is likely that the two morphotypes correspond to the botanical varieties *A. uniflora var. grandiflora* and *var. uniflora* (Gillet *et al.*, 1971). ITS and matK sequencing for the samples JFA_51 and JFA_71 (examples of plants with well-visible and small flowers, respectively) revealed consistent differences while flow cytometry measurements were relatively homogenious (Fig. 1b, Fig. S2, Table S2) (Chaintreuil *et al.*, 2016). In both morphotypes, numerous green nodules with a spherical shape and a narrow neck, running on the stem were visible (Fig. 3c).

*A. cristata* was found in pristine marshes and slow-flowing streams between Majunga and Mitsinjo where they often formed important stands of flooded plants. At sites where *A. cristata* plants were found *A*.*evenia* plants could be also present (Fig. 3d). *A. cristata* has retained little attention until it was shown to be one of the genome donors of *A. afraspera* and being a sister species of the Asian *A. aspera*, both species being profusely stem nodulated (Chaintreuil *et al.*, 2013, 2016; Devi., 2013a,b). Whereas the *A. cristata* specimen characterized by Chaintreuil *et al.* (2016) had densely-hairy stems, those found and collected in Madagascar were glabrous. Based on this characteristic, they may be tentatively associated to *A. cristata var. cristata* (hairy stems) and var. pubescens (glabrous stems) as decribed by Gillet *et al.* (1971). However, sequence and flow cytometry data additional to those obtained for sample JFA_34 are required to assess any genetic differentiation (Fig. 1b, Fig. S3, Table S2). In all sampling sites, *A. cristata* specimens cought attention very quickly due to the profuse nodulation all over the stem (Fig. 3d). These stem nodules were green and hemisperical with a broad base as described for the related *A. afraspera* species (Alazard and Duhoux, 1987).

## Stem nodulation in Sesbania rostrata

S. rostrata was observed in only two sampling sites in the region of Majunga. *S. rostrata* has been used a research model for nodulation due to its profuse stem nodulation and its ability to switch from classical nodulation to Lateral Root Base nodulation in flooded conditions (Capoen *et al*., 2009; Dreyfus and Dommergues, 1981). *S. rostrata* is the single species of its lineage in the *Sesbania* phylogeny (Furraggia *et al.*, 2020). However, *S. rostrata* specimens of Senegal and Madagascar were shown to be morphologically different and both hybridization and grafting experiments were less successful when interspecific (Ndiaye, 2005). For this reason, it has been proposed they could represent different subspecies of *S. rostrata. ITS* and *matK* sequencing of *S. rostrata* samples JFA_21 and JFA_45 also point to genetic differentiation when compared to an accession from Senegal, while flow cytometry analysis revealed similar genome sizes for the same accessions (Fig. 1c, Fig. S4, Table S2). In plants of both sampling sites, nodulation all along the stem was remarkable for its profusion and nodulation sites were typically distributed in vertical rows (Fig. 3e). These stem nodules were green, prominent, and had a constricted base.

## Conclusions and perspectives

In 2001, James *et al*. identified several stem-nodulated legumes in the Brazilian Pantanal wetlands. To broaden our knowledge on stem nodulating legumes, this research inspired us to explore wetlands in a geographically distinct tropical region, Madagascar. In Malagasy wetlands, a large occurrence and diversity of stem nodulation in *Aeschynomene* and *Sesbania* legumes was detected. While some of the species found are already well-known to form stem nodules (*A. elaphroxylon, A. evenia, A. indica 4x, A. schimperi, A. sensitiva and S. rostrata*), for two other identified species, *A. cristata* and *A. uniflora*, this has been once furtively mentioned in a review article and subject to very limited experimentations (Ladha *et al.*,1992; Chaintreuil *et al.*, 2016). Here, we report on their stem nodulation in the field. Following the traditional classification of stem-nodulating legumes, *A. cristata* was convincingly a *bona fide* profusely stem-nodulated species, equaling the stem nodulation level found for *A. afraspera* and *S. rostrata* (Boivin *et al.*, 1997a; Ladha *et al*., 1992). *A. uniflora* stem nodulation did not reach such level in the field. However, in greenhouse conditions *A. uniflora* stem nodulation has been shown to be exceptionally dense and to occur all along the stem (Chaintreuil *et al.*, 2016), indicating a profuse stem-nodulation capacity. The discovery of all these species and the demonstration of genetic diversity in several of them (*A. cristata*, *A. uniflora*, and *S. rostrata*) point out that more systematic studies including the collection of plants to evaluate species and ecotypes endowed with stem-nodulation are required.

What lessons could we learn from these studies ? On the plant side, profuse stem nodulation may be indicative of high nitrogen fixation activity, the presence of two stem-nodule morphologies (either hemispherical with a broad attachment to the stem or spherical with a narrow base) supports the existence of two developmental programs, and in both cases the photosynthetic activity in stem nodules (inferred from their green color) points to a physiology that likely differs from those of root nodules (discussed in Ladha *et al.*, 1992; Legocki and Szalay, 1984). On the bacterial side, while *A. caulinodans* has been isolated from *S. rostrata* stem nodules in both Senegal and Madagascar (Boivin *et al.*, 1997b), a great variety of *Aeschynomene-*nodulating *Bradyrhizobium* strains do exist but their genetics are insufficiently understood (Molouba *et al.*, 1999; Miché *et al.*, 2011; Okazaki *et al.*, 2016). New photosynthetic strains having or lacking *nod* genes are expected to be present in stem nodules of *A. cristata* (because it is a parent species of *A. afraspera*) and of the Nod-independent *Aeschynomene* species (e.g., *A. evenia, A. indica 4x, and A. sensitiva*) respectively. Intringingly, previously only non-photosynthetic strains were isolated from *A. uniflora* root nodules (Molouba *et al*., 1999). It would thus be very interesting to investigate the nature of those present in its stem nodules. Given all the valuable information that can be gained from stem-nodulating legumes, we advocate reviving their study as this would significantly increase our understanding on the diversity of mechanisms underlying the nitrogen-fixing symbiosis in legumes.

## Acknowledgements

We thank Robin Duponnois (LSTM Laboratory, IRD) for providing assistance in developping this research work, Bernard Dreyfus (LSTM Laboratory, IRD) for his helpful comments on Malagasy legumes, and the IRD Institute for funding Jean-François Arrighi’s mission to Madagascar. We are also thankful to Dr Eric Giraud (PHIM Laboratory, IRD) for his critical reading of this manuscript. The present work has benefited from the facilities and expertise of the cytometry facilities of Imagerie-Gif (https://www.i2bc.paris-saclay.fr/bioimaging/). This work was supported by the French National Research Agency (ANR-SymWay-20-CE20-0017-04).

## Author contributions

JFA and NN wrote the manuscript. FFM, MFR, CC, ATER, MDA, H Randriambanona, H Ramanankierana and JFA carried out specimen collection, photography, identification and herbarium conservation. FF, MP and MB performed sequence and flow cytometry analyses.

## Data availability

Data obtained in this study are listed in Tables S1 to S3, and the methods are given in Methods S1. The DNA sequences generated in this study were deposited in GenBank under accession numbers OR448903-OR448909 (nuclear *ITS*) and OR463925-OR463932 (chloroplastic *matK*).

## Methods S1 Material and methods

### Description of the collecting areas

To select collection sites of *Aeschynomene* and Sesbania species in Madagascar, we made use of general information about their distribution as described in the compendium “The Leguminosae of Madagascar” (Dupuy, 2002) and utilized precise location data of previous isolated specimens present in collections through the Global Biodiversity Information Facility (GBIF - https://www.gbif.org/) and the Tropicos database (https://www.tropicos.org/). This resulted in the definition of four collecting areas: 1) the region of Majunga including the vicinity of the Kinkony lake where many temporary to permanent marshes locally named « matsabory » are present. This area also comprises the plain of Marovoay where numerous ricefields form the main rice granary of Madagascar, 2) the Itasy region in the Center of Madagascar where the Itasy Lake and ricefields are present, 3) the Alaotra region in the East side of Madagascar that corresponds to a large basin containing the Alaotra Lake. The presence of extensive ricefields make this region the second rice granary of Madagascar, and 4) Nosy Be located towards the North of Madagascar where in the whole region ricefields are present. Expeditions to the four regions were made in April and May 2023, at the end of the rainy season that corresponds to the flowering period for most *Aeschynomene* and Sesbania species. Plants were collected in the above mentionned areas but also « en route » from ricefields, rivers and marshes present along the RN1, RN2 and RN4 national roads and secondary roads leading to the Itasy lake, the Alaotra lake and the region of Majunga, respectively.

### Plant and data collection

At each sampling location, the presence or receding of water in the aquatic ecosystem was recorded. The stem nodulation status of individuals for each species present in the population was examined and correlated to their positions relative to the flooding area. Whenever possible, three individuals of each species were chosen at random to determine their root nodulation status. Both stems and roots of these individuals were photographed *in situ*. Plant material was frequently collected for germplasm conservation and production of voucher specimens. The latters were deposited at the Herbarium of the CNARP Institute in Antananarivo (Madagascar).

### Plant culture

Seeds collected in the field were dried at 34°C for one week and used for plant cultivation when fresh material production was required. Seed scarification and plant growth in the greenhouse were performed as indicated in Arrighi *et al.* (2012).

### Gene sequencing and sequence analysis

Genomic DNA was isolated from fresh leaves using the CTAB extraction method. The nuclear ribosomal internal transcribed spacer region (ITS: ITS1-5.8S rDNA gene-ITS2) and the chloroplast *matK* gene were amplified and sequenced as published in Chaintreuil *et al.* (2016). Additionnal ITS and *matK* sequences were retrieved from Chaintreuil *et al.* (2016, 2018) and Brottier *et al.* (2018) for *Aeschynomene* species and from Farrugia *et al.* (2018) for *Sesbania* species. To analyse sequence variations, sequences were aligned using Multalin (http://multalin.toulouse.inra.fr/multalin/multalin.html). ML phylogenetic tree reconstructions were obtained by aligning nucleotide sequences with the MUSCLE program that is incorporated in the MEGA X (v10.1.8) software. Aligned sequences were further processed in MEGA X using the maximum likelihood approach and the Kimura 2-parameter model with a 1000x bootstrap (BS).

### Genome size estimation

Flow cytometry measurements were performed using fresh leaf material as described in Arrighi *et al.*, 2012. Genome size estimations were based on the measurements of three plants per accession using *Lycopersicum esculentum* (Solanaceae) cv « Roma » (2c = 1.99 pg) as the internal standard.

