## Supplemental Figure 1 for "Stem nodulation: diversity and occurrence in *Aeschynomene* and *Sesbania* legumes from wetlands of Madagascar"

### Slide 1
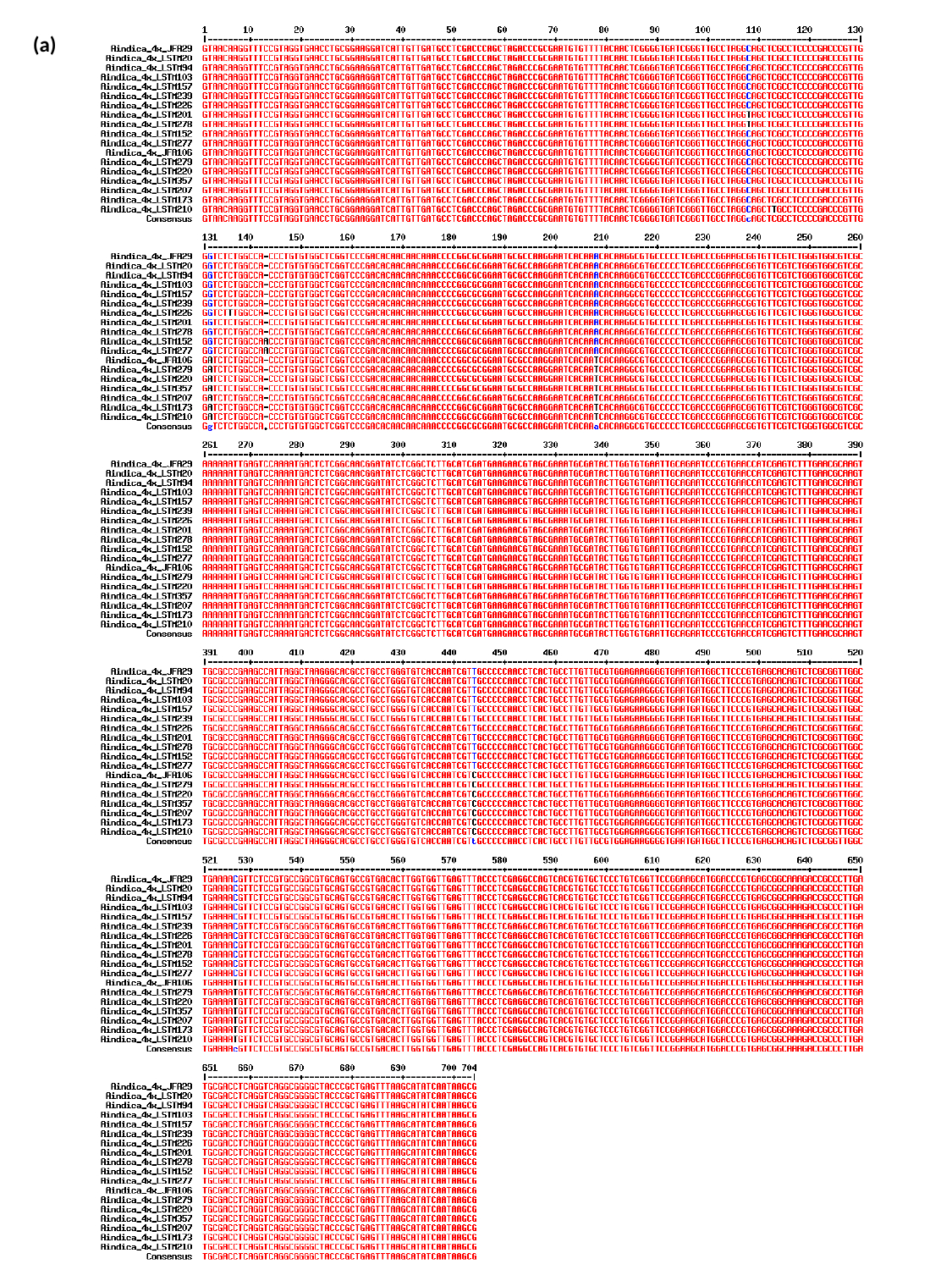

(a)

### Slide 2
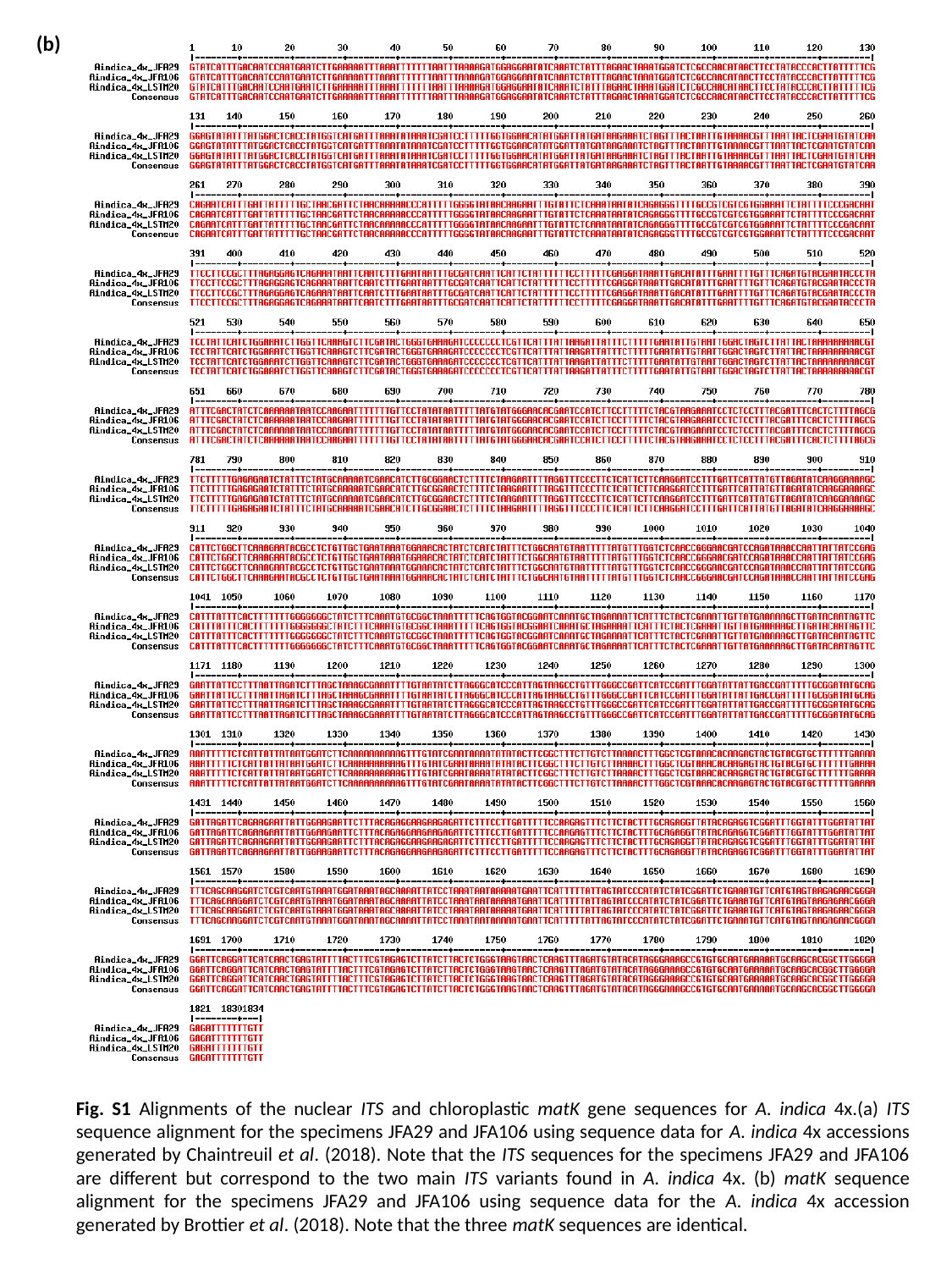

(b)
Fig. S1 Alignments of the nuclear ITS and chloroplastic matK gene sequences for A. indica 4x.(a) ITS sequence alignment for the specimens JFA29 and JFA106 using sequence data for A. indica 4x accessions generated by Chaintreuil et al. (2018). Note that the ITS sequences for the specimens JFA29 and JFA106 are different but correspond to the two main ITS variants found in A. indica 4x. (b) matK sequence alignment for the specimens JFA29 and JFA106 using sequence data for the A. indica 4x accession generated by Brottier et al. (2018). Note that the three matK sequences are identical.
