## Supplemental Figure 2 for "Stem nodulation: diversity and occurrence in *Aeschynomene* and *Sesbania* legumes from wetlands of Madagascar"

### Slide 1
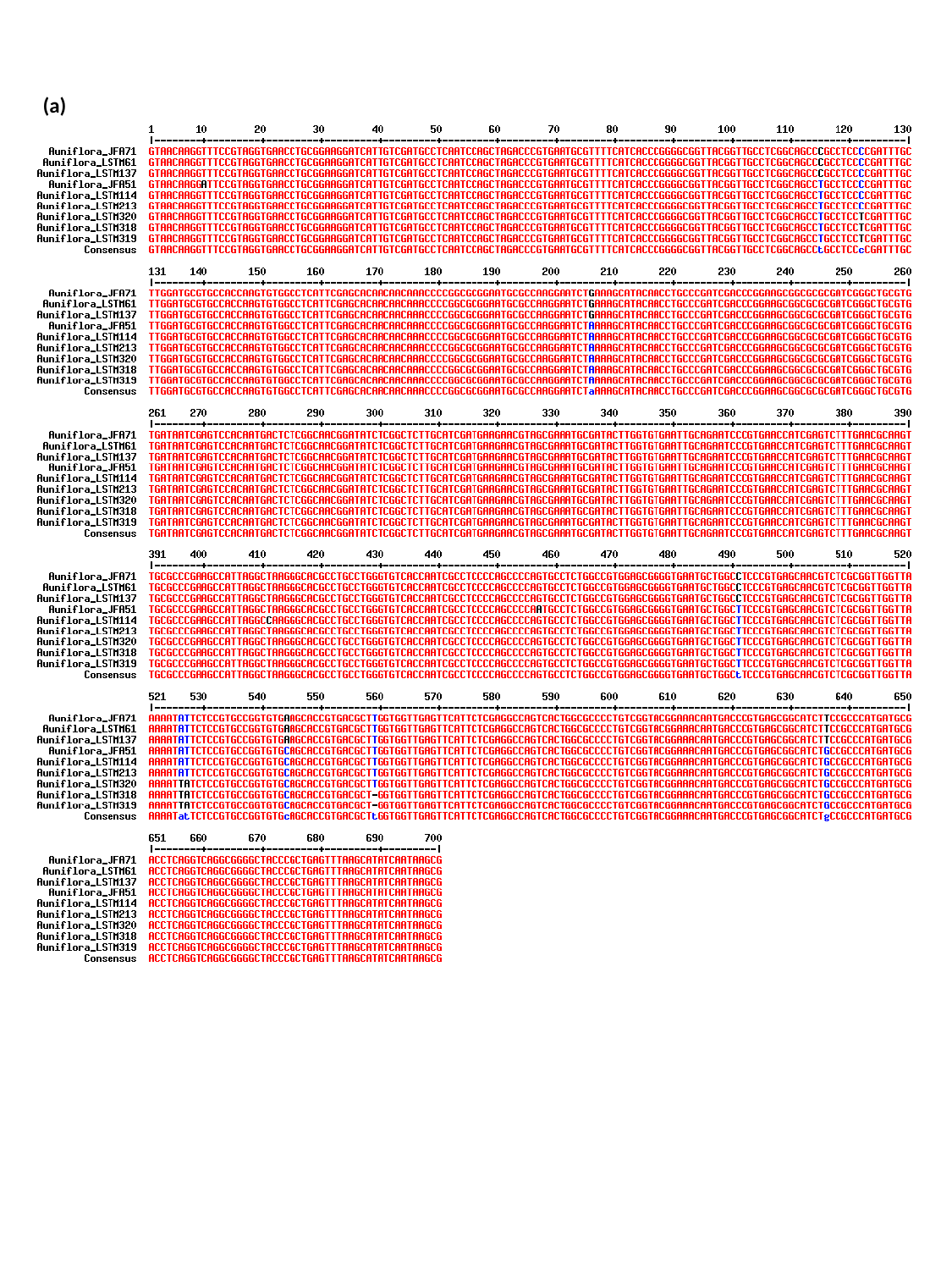

(a)

### Slide 2
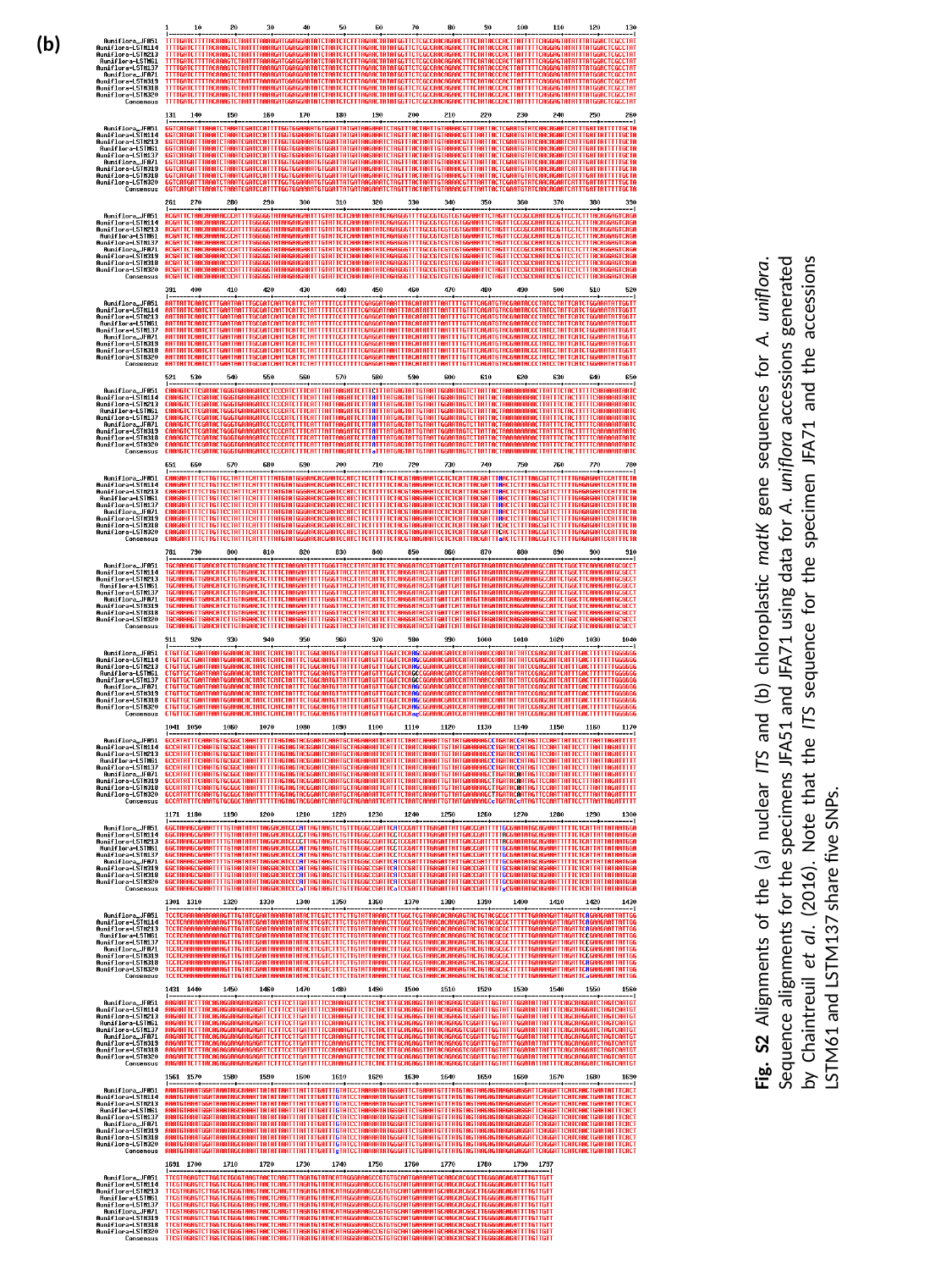

(b)
Fig. S2 Alignments of the (a) nuclear ITS and (b) chloroplastic matK gene sequences for A. uniflora. Sequence alignments for the specimens JFA51 and JFA71 using data for A. uniflora accessions generated by Chaintreuil et al. (2016). Note that the ITS sequence for the specimen JFA71 and the accessions LSTM61 and LSTM137 share five SNPs.
