## Supplemental Figure 3 for "Stem nodulation: diversity and occurrence in *Aeschynomene* and *Sesbania* legumes from wetlands of Madagascar"

### Slide 1
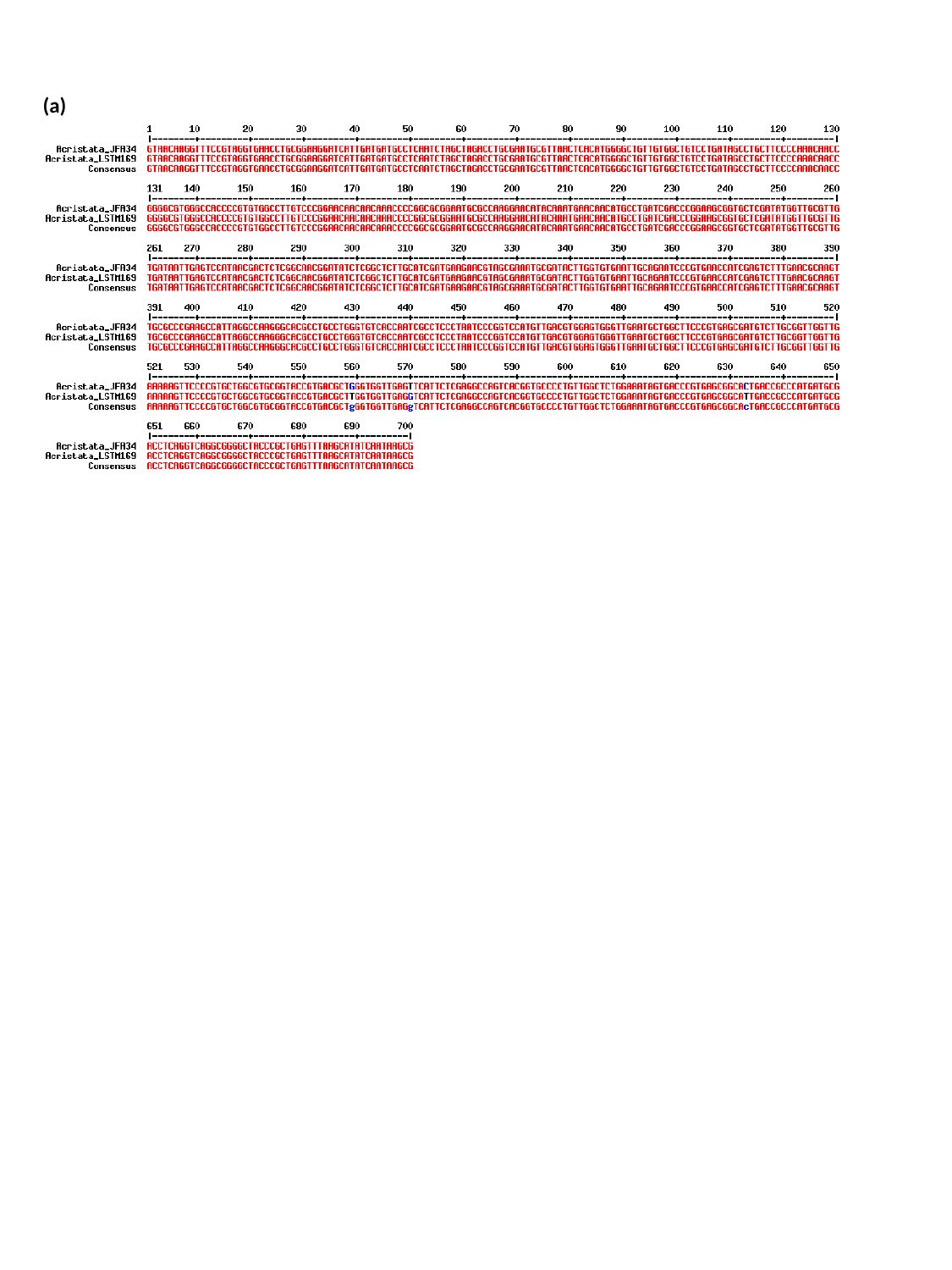

(a)

### Slide 2
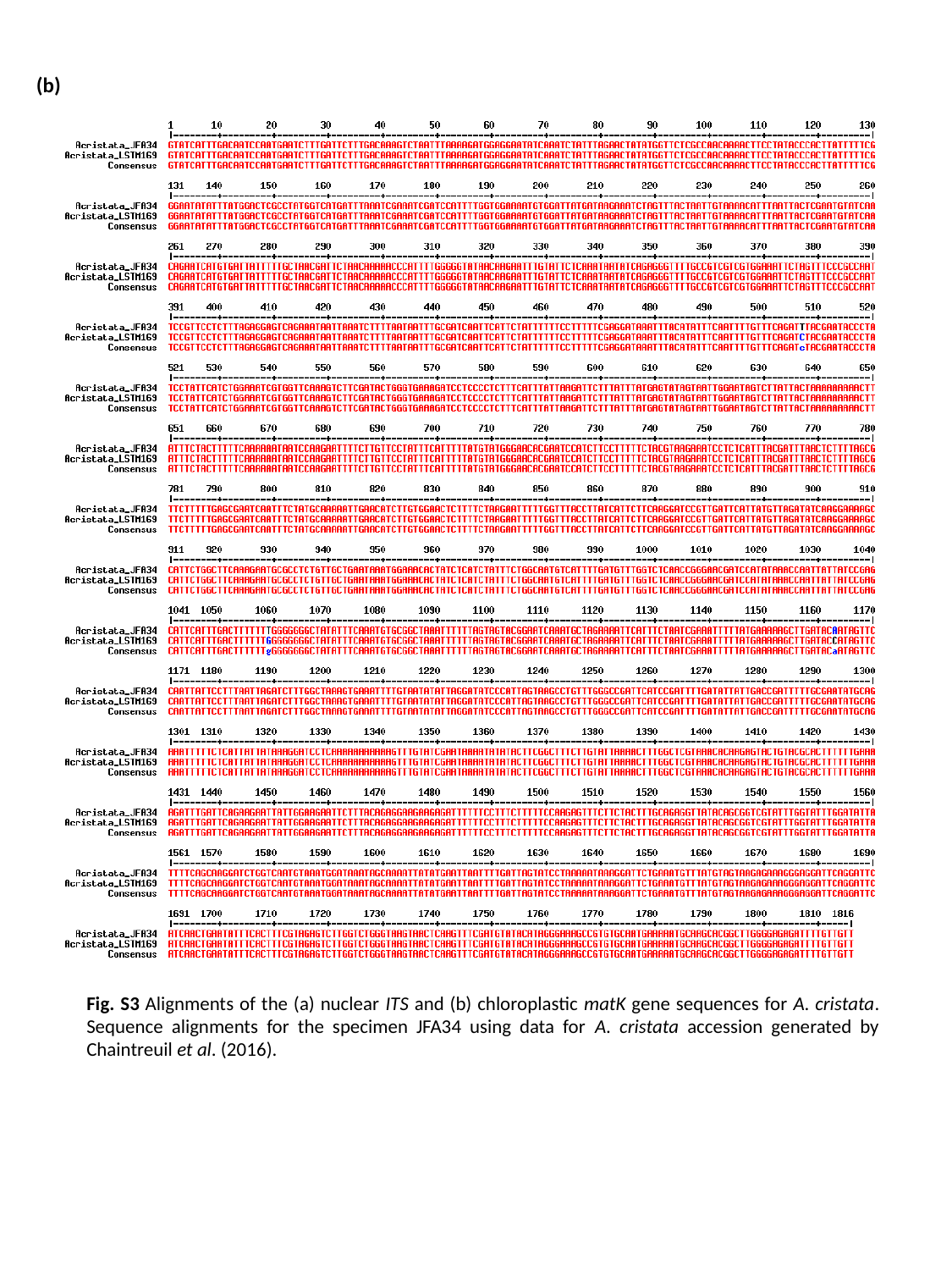

(b)
Fig. S3 Alignments of the (a) nuclear ITS and (b) chloroplastic matK gene sequences for A. cristata. Sequence alignments for the specimen JFA34 using data for A. cristata accession generated by Chaintreuil et al. (2016).
