## Supplemental Figure 4 for "Stem nodulation: diversity and occurrence in *Aeschynomene* and *Sesbania* legumes from wetlands of Madagascar"

### Slide 1
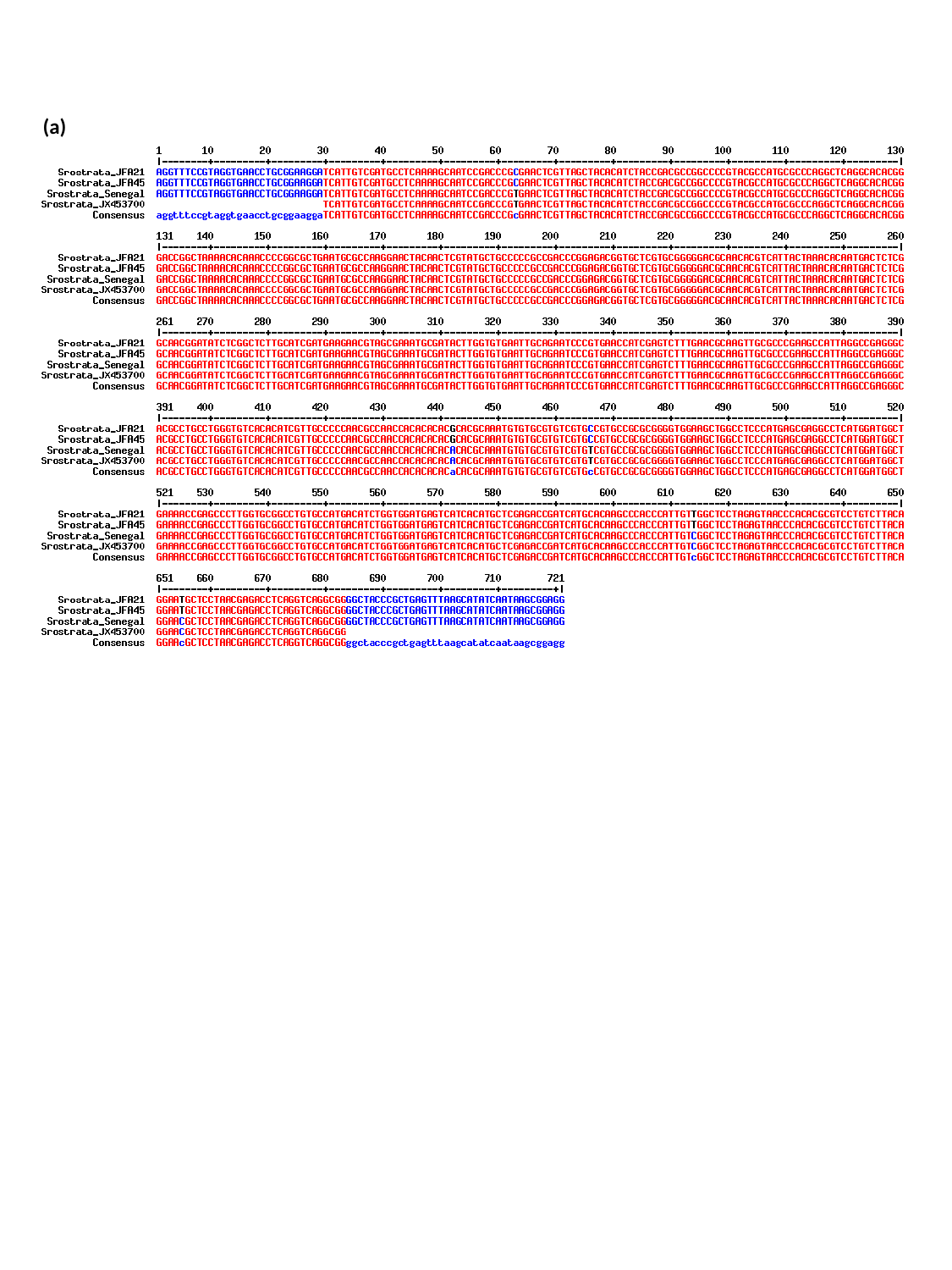

(a)

### Slide 2
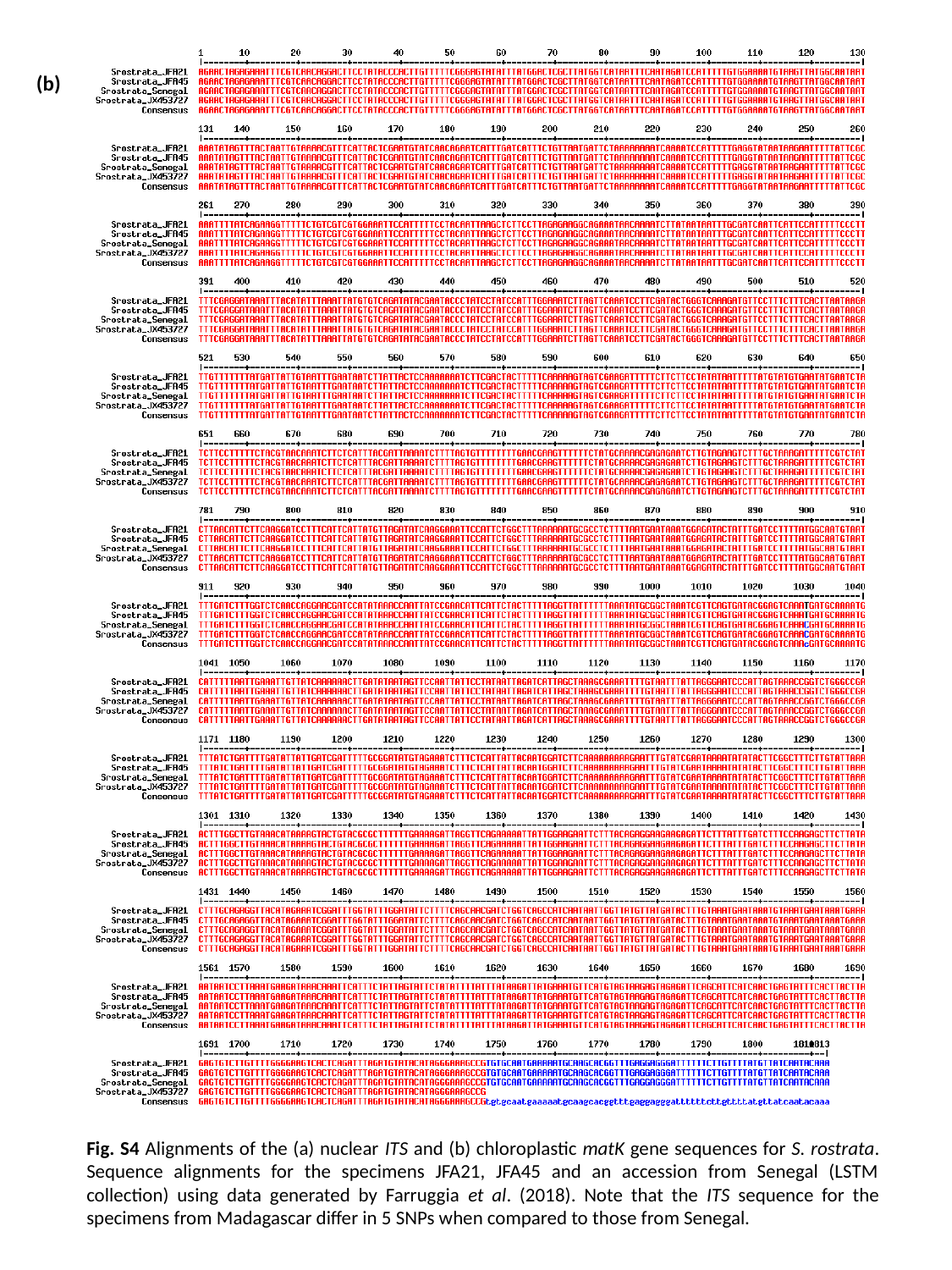

(b)
Fig. S4 Alignments of the (a) nuclear ITS and (b) chloroplastic matK gene sequences for S. rostrata. Sequence alignments for the specimens JFA21, JFA45 and an accession from Senegal (LSTM collection) using data generated by Farruggia et al. (2018). Note that the ITS sequence for the specimens from Madagascar differ in 5 SNPs when compared to those from Senegal.
