## Supplemental Table 1 for "Stem nodulation: diversity and occurrence in *Aeschynomene* and *Sesbania* legumes from wetlands of Madagascar"

| **Table S1** Accessions collected in this study, origin and characteristics | | | | |  |  |  |
| --- | --- | --- | --- | --- | --- | --- | --- |
| **Species** | **Collection code** | **Sampling area** | **Location** | **GPS coordinates** | **Situation** | **Nodules** | **CNARP voucher** |
| *A. cristata* | JFA34 | RN4-Majunga | Analamany | S 15° 52' 08.3" E 45° 52' 45.9" | Marsh | Stem | RFM167 |
|  | JFA39 | RN4-Majunga | Antangomenabevary | S 15° 58' 08.2" E 45° 59' 32.9" | Ricefield | Stem |  |
|  | JFA40 | RN4-Majunga | Besokatra | S15 °52' 25.1" E 46° 09' 03.1" | River | Stem | RFM165 |
|  | JFA41 | RN4-Majunga | Belemoka | S 15° 47' 59.5" E 46° 11' 07.1" | Marsh | Stem |  |
| *A. elaphroxylon* | JFA88 | RN2-Aloatra | Betoho | S 17° 46' 52.8" E 48° 26' 07.0" | Ricefield | Stem | RFM193 |
|  | JFA96 | RN2-Aloatra | Ambongabe | S 17° 44' 44.7" E 48° 26' 49.3" | Ricefield | Root | RFM201 |
| *A. evenia* | JFA2 | RN4-Majunga | Andohatapenaka | S 18° 54' 02.2" E 47° 29' 46.8" | Ricefield | Stem | RFM157 |
|  | JFA3 | RN4-Majunga | Mahitsy | S 18° 44' 22.0" E 47° 20' 21.2" | Ditch | Stem |  |
|  | JFA4 | RN4-Majunga | Anjomoka | S 18° 40' 26.3" E 47° 17' 23.7" | Ricefield | Stem |  |
|  | JFA15 | RN4-Majunga | Marofotroboka | S 16° 43' 11.2" E 47° 04' 05.3" | River | Stem | RFM163 |
|  | JFA17 | RN4-Majunga | Tsaramandroso | S 16° 21' 22.8" E 47° 01' 32.7" | Marsh | Stem |  |
|  | JFA23 | RN4-Majunga | Belobaka | S 15° 42' 27.0" E 46° 23' 29.5" | Marsh | Stem |  |
|  | JFA28 | RN4-Majunga | Bekopaka | S 15° 41' 45.0" E 46° 22' 33.5" | Marsh | Root |  |
|  | JFA33 | RN4-Majunga | Andrefamanangy | S 15° 57' 42.2" E 45° 57' 36.7" | Marsh | Stem | RFM166 |
|  | JFA35 | RN4-Majunga | Analamany | S 15° 52' 08.3" E 45° 52' 45.9" | Marsh | Stem |  |
|  | JFA37 | RN4-Majunga | Majunga - Besohaba marsh | S 15° 59' 26.8" E 45° 52' 34.6" | Ricefield | Stem |  |
|  | JFA47 | RN4-Majunga | Majunga - Bevovoka marsh | S 16° 06' 03.2" E 46° 40' 52.2" | Ricefield | Stem | RFM175 |
|  | JFA50 | RN4-Majunga | Bongomena | S 16°05' 11.4" E 46° 44' 38.2" | Ricefield | Stem | RFM178 |
|  | JFA54 | RN4-Majunga | Antimimalandy crossing | S 16° 23' 36.8" E 47° 05' 47.9" | Ricefield | Stem | RFM182 |
|  | JFA56 | RN4-Majunga | Carrière | S 16° 25' 38.0" E 47° 08' 07.4" | Marsh | Stem | RFM184 |
|  | JFA58 | RN4-Majunga | Ampasambazaha | S 16° 56' 27.8" E 46° 49' 52.3" | Ricefield | Stem | RFM185 |
|  | JFA59 | RN4-Majunga | Andriba | S 17° 36' 26.4" E 46° 55' 46.6" | Ricefield | Stem | RFM186 |
|  | JFA72 | RN1-Itasy | Ampefy | S 19° 03' 25.6" E 46° 44' 30.4" | Ricefield | Stem | JFA72 |
|  | JFA77 | RN1-Itasy | Virgin Island | S 19° 04' 00.0" E 46° 45' 48.9" | Ricefield | Stem | JFA77 |
|  | JFA86 | RN2-Aloatra | Manakambahiny | S 18° 54' 22.2" E 47° 58' 03.1" | Ricefield | Stem | RFM190 |
|  | JFA94 | RN2-Aloatra | Camp Bandro | S 17° 38° 25.7" E 48° 30' 18.2" | Ricefield | Stem | RFM199 |
|  | JFA98 | RN2-Aloatra | Ambalabako | S 17° 50' 55.1" E 48° 25' 09.8" | Ricefield | Stem | RFM203 |
|  | JFA100 | RN2-Aloatra | Vodiala | S 17° 53' 04.1" E 48° 15' 23.2" | Ricefield | Stem | RFM205 |
|  | JFA103 | RN2-Aloatra | Ambohimangakely | S 18° 53' 31.4" E 47° 36' 41.6" | Ricefield | Stem |  |
|  | JFA109 | Nosy Be | Nosy Be - Andilana | S 13° 15' 14.6" E 48° 11' 14.4" | Ricefield | Stem |  |
|  | JFA112 | Nosy Be | Nosy Be - West road | S 13° 22' 37.3" E 48° 13' 06.9" | Ricefield | Stem |  |
| *A. indica* 4x | JFA27 | RN4-Majunga | Majunga - Amparihingidro marsh | S 15° 42 '21.7" E 46° 23' 16.1" | Marsh | Stem | RFM170 |
|  | JFA29 | RN4-Majunga | Majunga - Petite Plage marsh | S 15° 39' 34.7" E 46° 20' 05.4" | Marsh | Stem | RFM169 |
|  | JFA106 | Nosy Be | Nosy Be - East road | S 13° 27' 46.1" E 48° 19' 16.1" | Ricefield | Stem |  |
|  | JFA107 | Nosy Be | Nosy Be - East road | S 13° 17' 20.0" E 48° 18' 38.5" | Ricefield | Stem |  |
| *A. schimperi* | JFA1 | RN1-Itasy | Antananarivo- Tsimbazaza park | S 18° 55' 46.2" E 47° 31' 31.7" | Garden | Root |  |
|  | JFA6 | RN4-Majunga | Anjomoka | S 18° 40' 26.3" E 47° 17' 23.7" | Ricefield | Root | RFM159 |
|  | JFA62 | RN1-Itasy | Maharefo | S 19° 01' 07.9" E 47° 11' 37.9" | Ricefield | Stem | JFA62 |
|  | JFA64 | RN1-Itasy | Ankadimena | S 19° 56' 47.4' E 49° 51' 45.4" | Ricefield | Stem | JFA64 |
|  | JFA68 | RN1-Itasy | Ankazomasina | S 18° 57' 11.6" E 46° 40' 06.0" | Ricefield | Stem | JFA68 |
|  | JFA70 | RN1-Itasy | Ampefy | S 19° 03' 25.6" E 46° 44' 30.4" | Ricefield | Stem | JFA70 |
|  | JFA74 | RN1-Itasy | Mahiatrondro | S 19° 01' 15.8" E 46° 43' 10.0" | Ricefield | Stem | JFA74 |
|  | JFA81 | RN2-Aloatra | Sambaina | S 18° 53' 28.7" E 47° 47' 20.5" | Ricefield | Stem |  |
|  | JFA84 | RN2-Aloatra | Carion | S 18° 54' 37.6" E 47° 42' 13.9" | Ricefield | Stem | RFM189 |
|  | JFA90 | RN2-Aloatra | Ambotrasana | S 17° 43' 10.3" E 48° 27' 24.7" | Ricefield | Stem | RFM195 |
|  | JFA97 | RN2-Aloatra | Ambalabako | S 17° 50' 55.1" E 48° 25' 09.8" | Ricefield | Stem | RFM202 |
|  | JFA102 | RN2-Aloatra | Ambohimangakely | S 18° 53' 31.4" E 47° 36' 41.6" | River | Root |  |
| *A. sensitiva* | JFA13 | RN4-Majunga | Betsiboka bridge | S 16° 56' 26.1" E 46° 57' 13.0" | Garden | Stem | RFM162 |
|  | JFA14 | RN4-Majunga | Marofotroboka | S 16° 43' 11.2" E 47° 04' 05.3" | River | Stem |  |
|  | JFA19 | RN4-Majunga | Amboronozy plains | S 16° 09' 40.0" E 46° 41' 02.8" | Marsh | Stem |  |
|  | JFA24 | RN4-Majunga | Majunga - Belobaka marsh | S 15°42' 27.0" E 46° 23' 29.5" | Marsh | Stem | RFM171 |
|  | JFA46 | RN4-Majunga | Mandikanamana | S 16° 06' 55.9" E 46° 43' 23.7" | Ricefield | Stem | RFM174 |
|  | JFA52 | RN4-Majunga | Bemailaka | S 16° 20' 38.4" E 46° 50' 40.0" | Marsh | Stem | RFM180 |
|  | JFA67 | RN1-Itasy | Ankadinondry | S 19° 00' 16.9' E 46° 27' 08.2" | Ricefield | Stem | JFA67 |
|  | JFA85 | RN2-Aloatra | Manakambahiny | S 18° 54' 22.2" E 47° 58' 03.1" | Ricefield | Stem | RFM191 |
|  | JFA87 | RN2-Aloatra | Moramanga | S 18° 56' 42.0" E 48° 13' 32.6" | Ricefield | Stem | RFM192 |
|  | JFA89 | RN2-Aloatra | Tanambao-biampasika | S 17° 45' 51.6" E 48° 26' 29.6" | River | Stem | RFM194 |
|  | JFA93 | RN2-Aloatra | Ambatosoratra | S 17° 36' 20.9" E 48° 30' 55.8" | Ricefield | Stem | RFM198 |
|  | JFA99 | RN2-Aloatra | Ambalavato | S 17° 51' 18.3" E 48° 18' 43.4" | Ricefield | Stem | RFM204 |
|  | JFA101 | RN2-Aloatra | Andranovelana | S 18° 17' 51.0" E 48° 16' 02.5" | Ricefield | Stem | RFM206 |
|  | JFA104 | Nosy Be | Nosy Be - South East | S 13° 22' 47.4" E 48° 19' 53.9" | Ricefield | Stem |  |
| *A. uniflora* | JFA51 | RN4-Majunga | Bongomena | S 16° 05' 11.4" E 46° 44' 38.2" | Ricefield | Stem | RFM179 |
|  | JFA53 | RN4-Majunga | Bemailaka | S 16° 20'38.4" E 46° 50' 40.0" | Marsh | Stem | RFM181 |
|  | JFA71 | RN1-Itasy | Ampefy | S 19° 03' 25.6" E 46° 44' 30.4" | Ricefield | Stem | JFA71 |
|  | JFA80 | RN1-Itasy | Soavinandriana | S 19° 11' 12.8" E 46° 45' 50.4" | Ricefield | Root | JFA80 |
|  | JFA108 | Nosy Be | Nosy Be - Ampasindava | S 13° 16' 17.4" E 48° 16' 51.8" | Ricefield | Stem |  |
|  | JFA111 | Nosy Be | Nosy Be - West road | S 13° 22' 37.3" E 48° 13' 06.9" | Ricefield | Stem |  |
| *S. rostrata* | JFA21 | RN4-Majunga | Amboromalandy | S 16° 07' 08.8" E 46° 45' 06.1" | Marsh | Stem | RFM164 |
|  | JFA45 | RN4-Majunga | Ambalanomby | S 16° 06' 28.3" E 46° 39' 16.4" | Ricefield | Stem |  |
