## Supplemental Table 2 for "Stem nodulation: diversity and occurrence in *Aeschynomene* and *Sesbania* legumes from wetlands of Madagascar"

| **Table S2** Nuclear 2C DNA amounts obtained by flow cytometry. | | | | |
| --- | --- | --- | --- | --- |
| **Species** | **Code** | **2C DNA content (pg)** | **Origin** | **Reference** |
| *A. cristata* | JFA34 | 1,81 ± 0.00 | Madagascar | This study |
|  | ILRI16880 | 1.92 ± 0.02 | CAR | Chaintreuil *et al*. (2016) |
| *A. indica 4x* | JFA29 | 1,72 ± 0.02 | Madagascar | This study |
|  | JFA106 | 1,67 ± 0.01 | Madagascar | This study |
|  | CIAT 17341 | 1.71 ± 0.07 | Thailand | Arrighi *et al*. (2014) |
| *A. uniflora* | JFA51 | 2.78 ± 0.02 | Madagascar | This study |
|  | JFA71 | 2.73 ± 0.01 | Madagascar | This study |
|  | LSTM319 | 2.70 ± 0.04 | Madagascar | Chaintreuil *et al.* (2016) |
| *S. rostrata* | JFA21 | 2.57 ± 0.02 | Madagascar | This study |
|  | JFA45 | 2.58 ± 0.02 | Madagascar | This study |
|  | Senegal | 2.59 ± 0.01 | Senegal | This study |
