## Supplemental Table 3 for "Stem nodulation: diversity and occurrence in *Aeschynomene* and *Sesbania* legumes from wetlands of Madagascar"

| **Table S3** GenBank numbers for the sequences generated in this study. | | | |
| --- | --- | --- | --- |
| **Species** | **Accession** | ***ITS*** | ***matK*** |
| *A. cristata* | JFA34 | OR448910 | OR463928 |
| *A. indica* 4x | JFA29 | OR448906 | OR463929 |
| *A. indica* 4x | JFA106 | OR448907 | OR463930 |
| *A. uniflora* | JFA51 | OR448908 | OR463931 |
| *A. uniflora* | JFA71 | OR448909 | OR463932 |
| *S. rostrata* | Senegal | OR448903 | OR463925 |
| *S. rostrata* | JFA21 | OR448904 | OR463926 |
| *S. rostrata* | JFA45 | OR448905 | OR463927 |
